# *Drosophila* p38 MAPK Interacts with BAG-3/starvin to Regulate Age-dependent Protein Homeostasis

**DOI:** 10.1101/552729

**Authors:** Sarah M. Ryan, Michael Almassey, Amelia M. Burch, Gia Ngo, Julia M. Martin, David Myers, Devin Compton, Scott Barbee, Nathan T. Mortimer, Subhabrata Sanyal, Alysia D. Vrailas-Mortimer

## Abstract

As organisms age, they often accumulate protein aggregates that are thought to be toxic, potentially leading to age-related diseases. This accumulation of protein aggregates is partially attributed to a failure to maintain protein homeostasis. A variety of genetic factors have been linked to longevity, but how these factors also contribute to protein homeostasis is not completely understood. In order to understand the relationship between aging and protein aggregation, we tested how a gene that regulates lifespan and age-dependent locomotor behaviors, p38 MAPK (p38Kb), influences protein homeostasis as an organism ages. We find that p38Kb regulates age-dependent protein aggregation through an interaction with the Chaperone-Assisted Selective Autophagy complex. Furthermore, we have identified Lamin as an age-dependent target of p38Kb and the Chaperone-Assisted Selective Autophagy complex.

## Introduction

Protein turnover is critical for maintaining tissue health as many proteins become damaged or misfolded during normal tissue functions. Therefore, the cell utilizes a variety of protein quality control mechanisms to refold or degrade these damaged proteins, including the ubiquitin proteasome system and macroautophagy. During aging, protein quality control mechanisms become less efficient leading to the accumulation of damaged or misfolded proteins that begin to form protein aggregates^1^. It has been hypothesized that these aggregates are toxic and may lead to the deleterious phenotypes associated with normal aging, such as impaired tissue function^1^. Furthermore, decreased protein aggregation has been associated with longevity. For example, over-expression of Foxo leads to an increased lifespan but also a concordant decrease in protein aggregation in *C. elegans*, *Drosophila,* and mice ^2–6^, suggesting that lifespan and protein aggregation are tightly linked processes. However, the molecular mechanisms that underlie the relationship between aging and protein homeostasis have not been fully characterized.

One pathway that has been linked to both aging and protein homeostasis is the stress response p38 MAPK (p38K) pathway. In mammals, there are four p38K genes (α, β, γ, and δ), and p38Kα has been linked to both the inhibition ^7,8^ and induction ^9,10^ of macroautophagy, in particular in response to oxidative stress ^11,12^. In addition, p38Kα has been linked to regulating macroautophagy in cellular senescence ^13–15^. However, how p38K signaling may contribute to protein homeostasis in response to natural aging is not well understood. The fruit fly *Drosophila melanogaster* has two canonical p38K genes (p38Ka and p38Kb), and we have previously reported that p38Kb acts in the adult musculature to regulate aging. We found that over-expression of p38Kb leads to increased lifespan while loss of p38Kb results in a short lifespan and age-dependent locomotor behavior defects ^16^. In addition, oxidatively damaged proteins accumulate in the muscle of p38Kb mutants with age ^16^, and loss of p38Kb leads to increased polyubiquitination of insoluble proteins and alterations in oxidative stress dependent translation ^17^, suggesting that these oxidatively damaged proteins may be aggregating in p38Kb mutants. Furthermore, p38Kb has been shown in a *Drosophila* cell culture system to pull down with the HspB8 homologue CG14207 ^18^, which plays a role in the muscle by regulating protein homeostasis as a part of a protein quality control mechanism called BAG-3 Mediated Selective Macroautophagy pathway in mammalian systems or the Chaperone-Assisted Selective Autophagy (CASA) complex in flies ^19,20^. The CASA complex also includes the chaperone Hsc70 and the co-chaperone BAG-3 (starvin in flies). BAG-3/starvin binds to specific damaged or misfolded protein substrates and brings them to the CASA complex where it binds to Hsc70 and HspB8. Those substrates that cannot be refolded by the complex are polyubiquitinated and targeted to the autophagosome through a handover between BAG-3/stv and the autophagy adaptor protein p62 (ref(2)p in flies), and subsequently degraded through the autophagosome-lysosome system ^21–25^.

Here, we report that p38Kb regulates age-dependent muscle protein homeostasis through an interaction with the CASA complex. We find that p38Kb acts as an intermediary between BAG-3/starvin and p62/ref(2)p in targeting damaged or misfolded proteins for degradation. This interaction is not only important for maintaining protein homeostasis but also for lifespan extension. In addition, we find that the *Drosophila* homologue of the Hutchinson-Gilford progeria protein, Lamin A/C, is a target for p38Kb and CASA complex mediated protein turnover, suggesting that the p38Kb aging phenotypes may be a result of impaired Lamin degradation.

## Materials and Methods

### Genotypes

UAS-p38Kb wt, UAS-p38Kb Kinase Dead, p38Kb ^Δ45^, p38Kb^Ex41^, w^1118^, Mef2-GAL4 and MHC-GAL4 were as described in ^16^. The p38Kb^Ex41^ is a precise excision allele and serves as a genetic background control for p38Kb ^Δ45^ deletion mutation.

UAS-stv RNAi 34408 (w^1118^; P{GD10796}v34408) and UAS-stv RNAi 34409 (w^1118^; P{GD10796}v34409/TM3) are described in ^26^ and are from the Vienna Drosophila Resource Center.

stv-GFP trap (w^11118^;; Pbac{754.P.F3v30} stv^+CPTI^ ^002824^), HspB8-GFP trap (w^1118^ PBac{810.P.FSVS-2}CG14207^CPTI004445^), Hsc70-4 GFP trap (*w^1118^ PBac{544.SVS-1}N^CPTI002347^*) are from the Kyoto Stock Center.

w^1118^; P{y[+mDint2] w[BR.E.BR]=SUPor-P}ref(2)P^KG00926^, w^1118^;;+mDint2 EY4969 stvEP, Lam^A25^ pr^1^, P{UAS-mito-HA-GFP.AP}3, w[126]; P{w[+mC]=UAS-Hsc70-4.WT}B, and w^1118^; P{w[+mC]=UAS-Lam.GFP}3-3 were obtained from the Bloomington Drosophila Stock Center.

All fly stocks were backcrossed into the w^1118^ background and isogenized for 10 generations. All stocks were reared at 25°C in a 12hr:12hr light:dark cycle on standard fly food media.

### Immunofluorescence

Adult flies were fixed in 4% paraformaldehyde for 48hrs at 4°C. Indirect flight muscles were dissected in 1X PBS, permeabilized in 1X PBS 0.15% Triton-X 100, and blocked in NGS + 0.15% Triton-X 100. Samples were incubated in primary antibody at 4°C overnight, washed in 1X PBS 0.15% Triton-X 100, and incubated in secondary antibody at room temperature for 2hrs. Samples were mounted in Vectashield mounting medium (Vectorlabs) and visualized using a laser scanning confocal microscope. Antibodies: rabbit anti-GFP 1:400 (Invitrogen), mouse anti-FLAG M2 1:1000 (Sigma), rabbit anti-stv 1:1000 (gift of Jög Höhfeld), rat anti-α actinin 1:100 (Abcam), rabbit anti-ubiquitin linkage-specific K63 anti-mouse 1:200 (Abcam), IgG-Alexa Fluor 488 1:200 (Life Technologies), anti-mouse IgG-Alexa Fluor 568 1:500 (Life Technologies), anti-rabbit IgG-Alexa Fluor 488 1:500 (Life Technologies) and Rhodamine Phalloidin 1:2000.

### Protein aggregate analysis

Indirect flight muscles were prepared as described above from 9 individual flies per genotype. Protein aggregates were identified using mouse anti-polyubiquitin 1:1000 (Enzo Life Sciences). Three muscles from each individual fly were imaged as z-series and flattened into a single image as a max projection using confocal microscopy for a total of 27 muscles per genotype. Images were analyzed using Image J “Analyze Particles” function with a diameter of 100 pixels set for the minimum aggregate size. Aggregate number and size were analyzed using ANOVA followed by Tukey’s HSD using the R package “multcomp” to generate significance groups with each letter group being significantly different with a p value of ≤ 0.05. Within genotype/across time point analyses were performed using the Welch two sample t-test in R.

### Lifespan

For lifespan experiments using the UAS-p38Kb^wt^, ref(2)p −/+, p38Kb^Δ45/Δ45^, the stv EP (stv^wt^) lines, and their respective controls, virgin females were kept on standard molasses *Drosophila* media. Due to a change in lab food, the stv RNAi 34408 and stv RNAi 34409 lifespan experiments (with their respective controls) were performed on the standard Bloomington *Drosophila* media. Virgin flies were collected and reared at 25°C in a 12hour:12hour light:dark cycle. Flies were put on new food twice a week. The number of dead animals was scored daily. Lifespan was analyzed using a log rank test to compare genotypes with censored data on all genotypes and then on all pairwise comparisons using the R package “survival” with Benjamini and Hochberg correction (false discovery rate < 0.05).

### Co-immunoprecipitation

Flies were aged 1 week or 5 weeks. Forty thoraxes per genotype per condition were homogenized in high salt buffer (0.5 M KCl, 35% glycerol,10 mM HEPES pH 7.0, 5 mM MgCl2, 0.5 mM EDTA pH 8.0, 0.1% NP40 25 mM NaF, 1 mM Na2VO4, 1 mM DTT, Complete protease inhibitor). The lysate was flash frozen in liquid nitrogen and quickly thawed at 37°C. Then lysates were rocked at 4°C for 30 minutes and centrifuged at 14,200 x g for 30 minutes at 4°C. The supernatant was transferred to equilibrated beads anti-Flag (M2) agarose (Sigma) or anti-GFP agarose (Chromotek) and rocked for 2 hours at 4°C. Beads were collected using a magnetic bar and washed four times with IP buffer (50 mM HEPES pH 7.0, 100 mM KCl, 0.4% NP40 1.5 mM MgCl2, 5% glycerol, 25 mM Na, 1 mM Na2VO4, 1 mM EDTA, 1 mM DTT, Complete protease inhibitor). Lysates were then analyzed by immunoblotting using rabbit anti-GFP 1:1000 (Invitrogen), mouse anti-FLAG M2 1:1000 (Sigma), rabbit anti-phospho-p38 1:1000 (Cell Signaling Technologies), goat anti total p38 1:1000 (Santa Cruz Biotechnology), rabbit anti-stv 1:10,000 (gift of Jrög Höhfeld) or mouse anti-Lamin 1:1000 (DHSB).

### Immunoblotting

Wild type flies (w^1118^) were aged either for 3, 15, 30, and 45 days or for 1-5 weeks. Three thoraxes were dissected and homogenized in 1x Laemmli buffer. Immunoblots were performed as described in ^16^. Membranes were developed using SuperSignal West Femto kit (ThermoFisher) or Pierce ECL (ThermoFisher) and exposed on autoradiography film. Antibodies used were: rabbit anti-GFP 1:1000 (Invitrogen), rabbit anti-starvin 1:10,000 (gift of Jrög Höhfeld), mouse anti-actin 1:5,000,000 (Sigma), mouse anti-FLAG M2 (Sigma), rabbit anti-alpha tubulin (Cell Signaling Technologies), mouse anti-Lamin 1:100 (DHSB), rabbit anti-phospho Lamin A Ser22 1:1000 (Thermofisher), mouse anti-beta tubulin (E-10) 1:5000 (Santa Cruz Biotechnology), mouse anti-HRP 1:20,000 (Jackson Labs), rabbit anti-HRP 1:40,000 (Jackson Labs). Densitometry was performed using a minimum of three independent blots. For statistical analysis of protein expression level, pixel density of the tested protein was normalized within sample to the loading control. These values were then normalized to control to calculate fold change. The fold change values were analyzed by Student’s t-test or ANOVA ^27^ as appropriate.

### Sucrose Gradient Fractionation

p38Kb^Ex41/Ex41^ and p38Kb^Δ45/Δ45^ flies were aged three weeks. 30 thoraxes per genotype were dissected and homogenized in NP40 lysis buffer. Samples were centrifuged at 800xg for 10 minutes at 4°C. The supernatant was then transferred to a 15-50% sucrose discontinuous gradient. Samples were then ultracentrifuged at 55,000 rpm (201,000xg at rav) for 20 hours at 4°C in a TLS-55 in a Beckman Coulter Optima TLX Ultracentrifuge. 200μl fractions were collected and the pellet was resuspended in an equal volume of NP40 Lysis Buffer.

### Stv localization

Immunohistochemistry on p38Kb^Ex41/Ex41^ and p38Kb^Δ45/Δ45^ Indirect flight muscles was performed as described above. Confocal images from five individual flies per genoytpe were analyzed for average pixel density using ImageJ in three different non-overlapping locations on each muscle for a total of 15 measurements per genotype. Average pixel density was analyzed by Student’s t-test using R.

## Results

### p38Kb regulates age-dependent protein homeostasis

p38Kb null mutant animals (p38Kb^Δ45/Δ45^, a deletion of the p38Kb gene) exhibit age-dependent locomotor behavior defects and have a 48% lifespan reduction. In addition, biochemical analysis suggests that p38Kb null mutants have increased levels of insoluble polyubiquitinated proteins in the thoracic musculature of aged animals as observed by immunoblot analysis ^17^. However, protein aggregate size and distribution have not been visualized in the p38Kb mutants, nor is it known whether augmenting p38Kb activity in muscles leads to a change in protein homeostasis. Therefore, we analyzed how p38Kb expression influences protein aggregation. We find that p38Kb null mutants have an increased number of protein aggregates in the adult indirect flight muscle at 1 week and 3 weeks of age (Figure 1A-B and E-F, Table S1) and increased aggregate size with age (Figure 1G-H, Table S1) as compared to a genetic background control (p38Kb^Ex41/Ex41^, a precise excision allele). Furthermore, transgenic inhibition of p38Kb in the muscle using a dominant negative kinase dead construct (p38Kb^KD^, ^16^) also results in a significant increase in aggregate number, however aggregate size was not affected (Figure 1I-L and Table S2). Conversely, as both strong and moderate levels of p38Kb over-expression lead to an increased lifespan (37% and 14% extension, respectively) ^16^, we tested if p38Kb over-expression also affects protein aggregation. We find that both strong over-expression of p38Kb in the adult muscle using the Mef2-GAL4 driver (Figure 1C-D and M-P, Table S3) and moderate over-expression of p38Kb using the MHC-GAL4 driver (Figure 1Q-T and Table S4) leads to decreased protein aggregate number and size throughout the lifespan. It has been hypothesized that protein aggregate accumulation and increased size may be toxic, potentially explaining the decreased lifespan and age-dependent locomotor abnormalities in p38Kb^Δ45^ null mutants and the increased longevity in the p38Kb over-expression animals.

**Fig 1.**
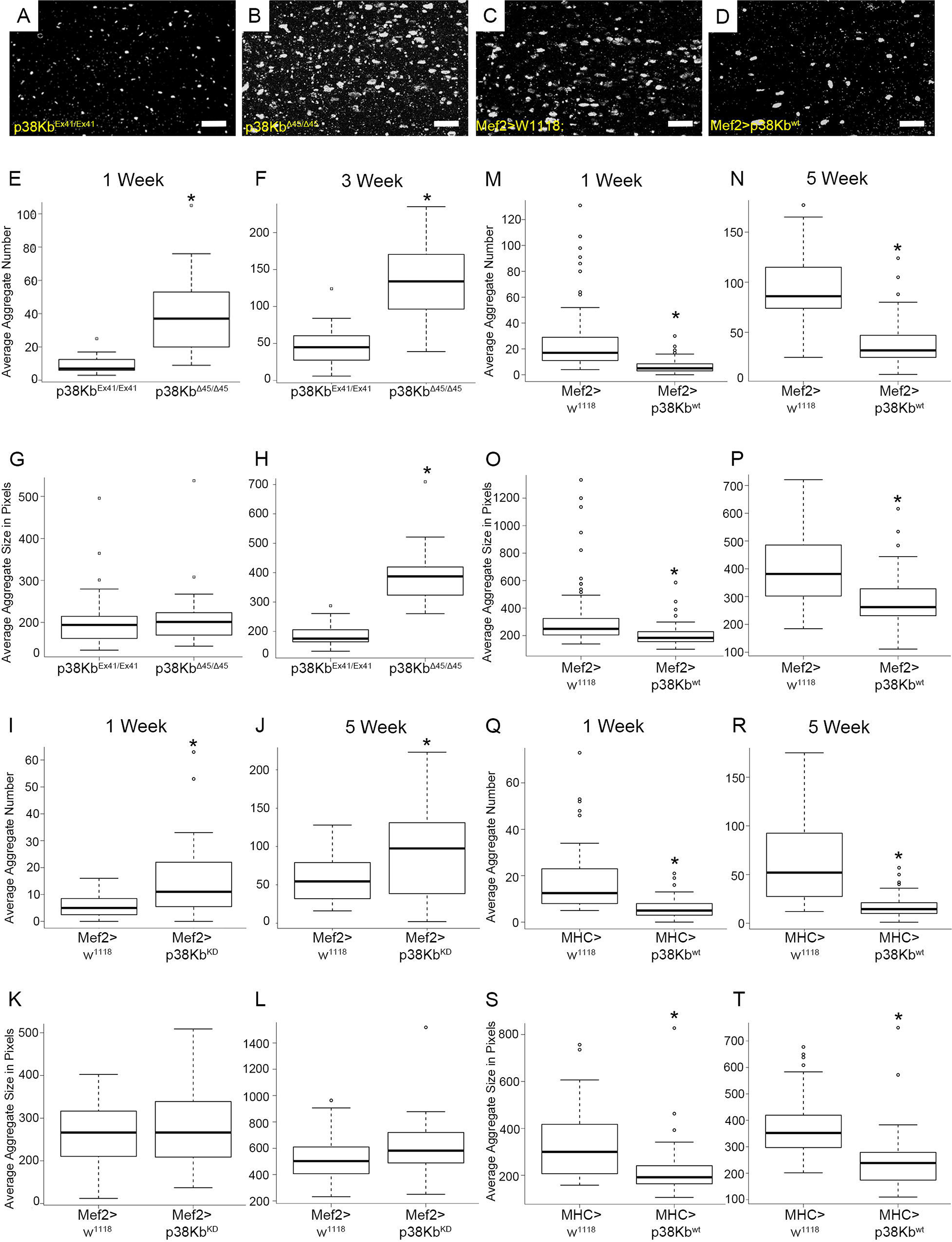
p38Kb regulates age-dependent protein homeostasis. Confocal micrographs of polyubiquitin positive protein aggregates in the adult indirect flight muscle in **A)** a precise excision genetic background control p38Kb^Ex41/Ex41^ and **B)** p38Kb ^Δ45/Δ45^ null mutants at three weeks of age and in **C)** outcrossed Mef2-GAL4 controls (Mef2>w^11118^) and **D)** UAS-p38Kb^wt^ Mef2-GAL4 (Mef2>p38Kb^wt^) over-expression animals at 5 weeks of age. Scale bar equals 6.2 μm. Box-Whisker plots of aggregate number in p38Kb ^Δ45/Δ45^ mutants as compared to p38Kb^Ex41/Ex41^ controls at **E)**1 week and **F)** 3 weeks of age and aggregate size at **G)**1 week and **H)**3 weeks of age. Aggregate number in p38Kb^KD^ Mef2-GAL4 (Mef2>p38Kb^KD^) and outcrossed Mef2-GAL4 controls at **I)** 1 week and **J)**5 weeks of age and aggregate size at **K)**1 week and **L)**5 weeks of age. Aggregate number in strong p38Kb over-expression animals and outcrossed Mef2-GAL4 controls at **M)**1 week and **N)**5 weeks of age and aggregate size at **O)**1 week and **P)**5 weeks of age. Aggregate number in moderate p38Kb over-expression (MHC>p38Kb^wt^) animals and outcrossed MHC-GAL4 controls (MHC>w^1118^) at **Q)**1 week and **R)**5 weeks of age and aggregate size at **S)**1 week and **T)**5 weeks of age.

### p38Kb mediates age-dependent phenotypes through autophagy

In order to determine what mechanism plays a role in the clearance of polyubiquitinated protein aggregates, we first tested what type of ubiquitin linkage is present in the aggregates in wild type flies. In particular, K63-linked ubiquitination has been shown to facilitate the formation of aggregates ^28–30^ that are cleared through autophagy ^30–32^. We find that a subset of aggregates from both aged wild type and p38Kb mutant flies contain K63-linked ubiquitinated proteins (Figure 2), suggesting that these muscle protein aggregates are degraded through autophagy.

**Fig 2.**
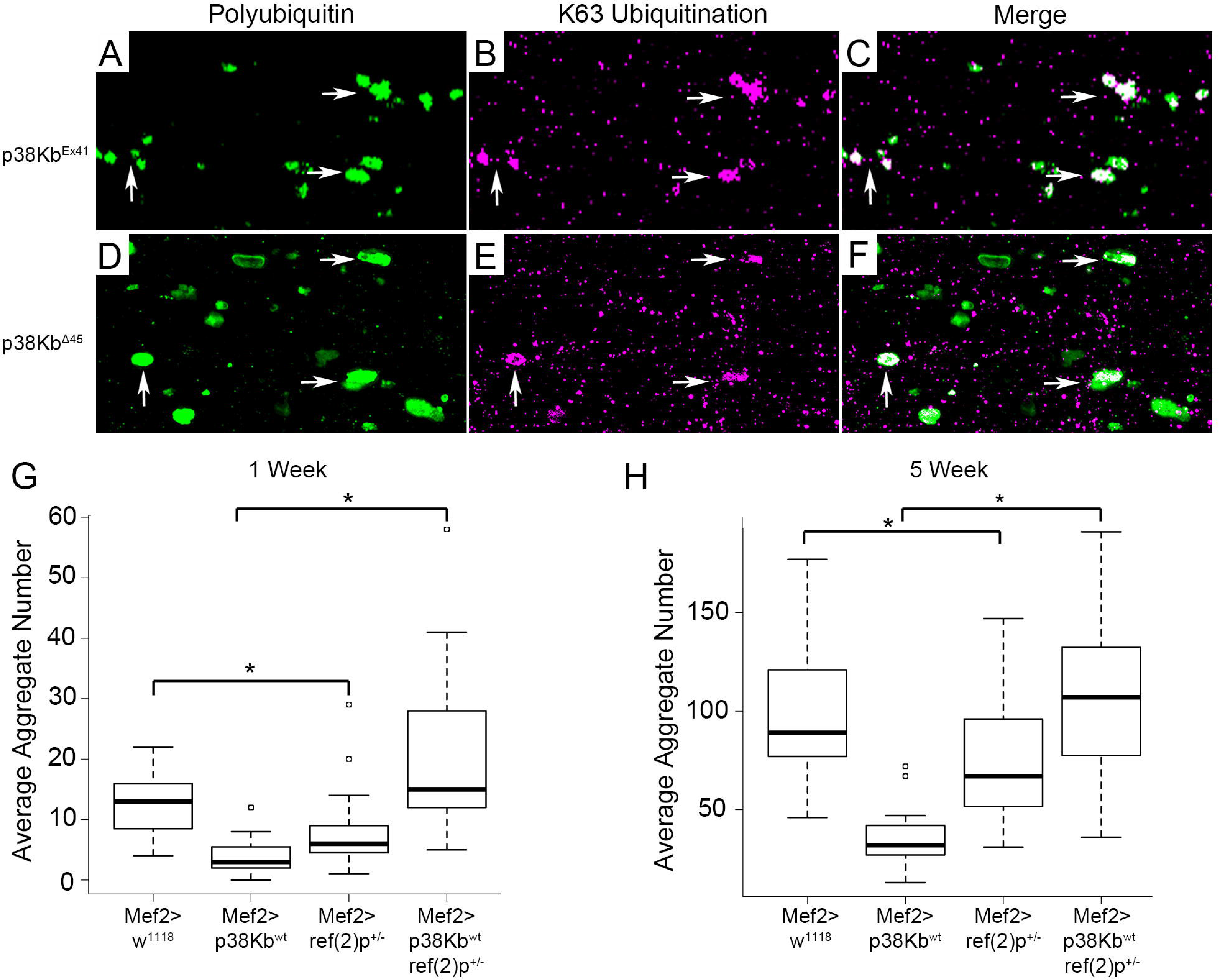
p38Kb regulates aging phenotypes through ref(2)p. **A-C)** A subset of polyubiquitinated protein aggregates (green) in 3 week old p38Kb^Ex4^/^Ex41^ control muscle contain K63 ubiquitinated (magenta) proteins (white arrows). **D-F)** K63 ubiquitin positive protein aggregates are also observed in 3 week old p38Kb^Δ45/Δ45^ mutant muscle. Protein aggregate number in ref(2)p heterozygous mutant backgrounds at **G)**1 week and **H)**5 weeks. Loss of a single copy of ref(2)p prevents the p38Kb mediated reduced protein aggregation at 1 week and 5 weeks of age. Asterisks denote a p-value of ≤0.001.

Polyubiquitinated protein aggregates can be degraded through selective autophagy in which the adaptor protein p62 (ref(2)p in flies) promotes the packaging and delivery of polyubiquitinated proteins to the autophagosome^33^. If p38Kb requires selective autophagy to mediate protein homeostasis, then loss of ref(2)p will block the p38Kb mediated decreased aggregation phenotype. We find that loss of a single copy of *ref(2)p* results in fewer aggregates (Figure 2G-H and Table S5), which may reflect compensation by other protein clearance mechanisms with aging, especially as homozygous *ref(2)p* mutants are viable. When a single copy of *ref(2)p* is removed in the p38Kb over-expression background, this prevents the reduced protein aggregation observed in the p38Kb over-expression animals at both young and old ages (Figure 2G-H and Table S5). This dominant interaction suggests that *ref(2)p* acts downstream of p38Kb to promote the degradation of protein aggregates.

### p38Kb colocalizes with the CASA complex in the adult flight muscle

In order to better understand the role of p38Kb in protein homeostasis, we first determined where in the muscle p38Kb localizes and find that a FLAG-tagged p38Kb (green) colocalizes with the Z-disc marker alpha-actinin and is also present at the M-line (Figure 3A). The muscle Z-disc is an area of high protein turnover, and one protein quality control mechanism that localizes to the Z-disc in mice and adult flies is the Chaperone-Assisted Selective Autophagy (CASA) complex (known as the BAG-3 mediated selective autophagy complex in mammals)^19,34,35^. In adult *Drosophila* muscle, the CASA complex has been reported to consist of the Hsc70 homologue Hsc70-4, the HspB8 homologue CG14207 and the BAG-3 homologue starvin (stv) ^19^. We find that p38Kb colocalizes with all the CASA complex members at the Z-disc and M-line (Figure 3B-D).

**Fig 3.**
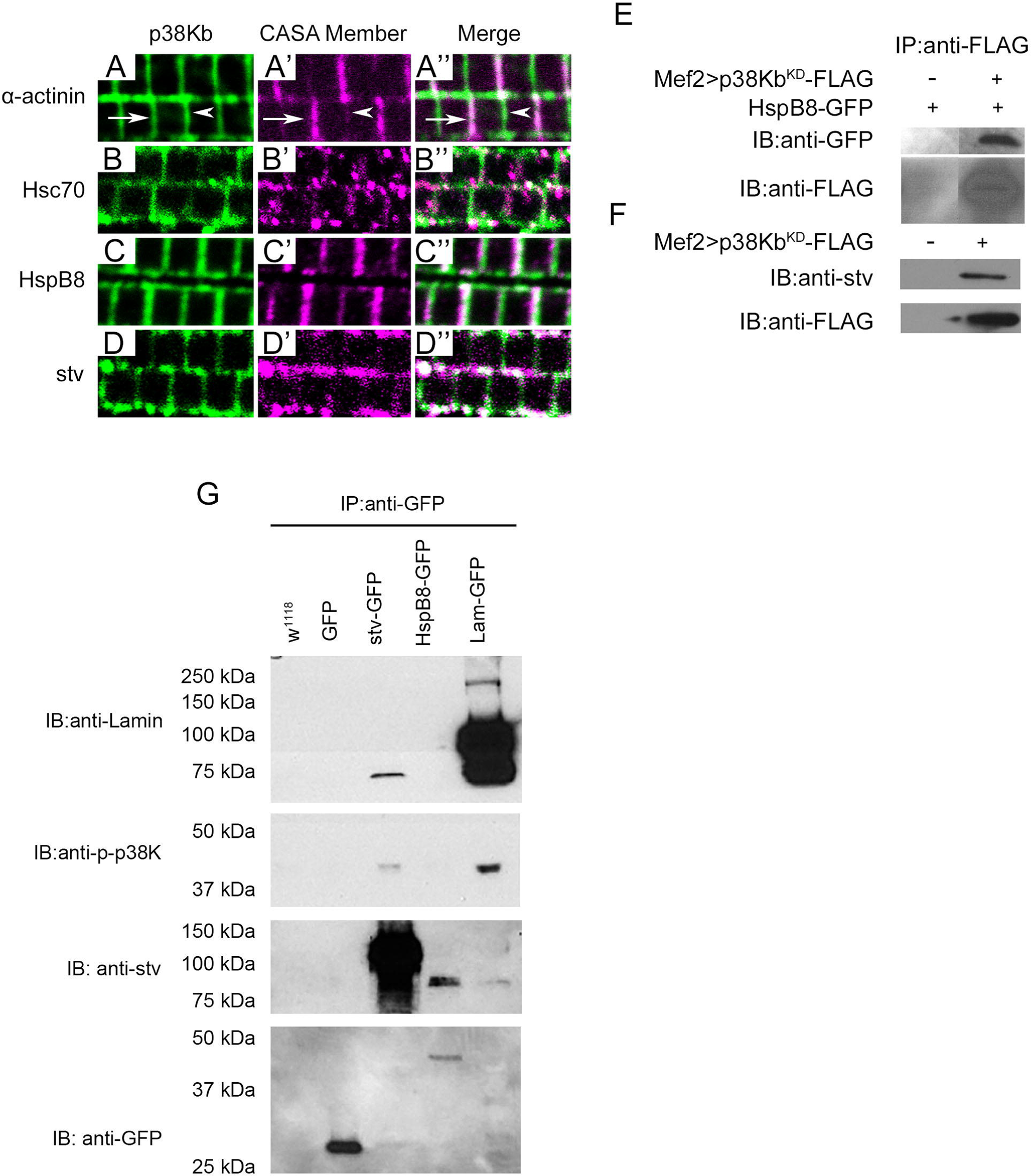
p38Kb genetically interacts with stv to regulate protein homeostasis and lifespan. **(A-D)** Localization of a FLAG-tagged p38Kb (green in **A-D** and **A’’-D’’**) in the adult indirect flight muscle. **A)** FLAG-tagged p38Kb localizes to the Z-disc (arrows) as exhibited by colocalization with the Z-disc protein alpha-actinin (magenta, **A’** and **A’’**), as well as the M-line (arrowheads). **B-D)** p38Kb colocalizes with each CASA complex member (magenta, **B’-D’**) at the Z-disc. Over-expression of a FLAG tagged p38Kb in the muscle in **B**) an endogenous Hsc70-GFP fusion protein background, **C**) an endogenous HspB8-GFP fusion protein background, and **D)** a wildtype background with endogenous levels of stv. **E-F)** Muscle lysates expressing control and Mef2-GAL4 UAS-p38Kb^KD-FLAG^ in either **E)** an endogenously expressed HspB8-GFP background or **F)** a wildtype background were immunoprecipitated using anti-FLAG coated beads. **E)** Immunoblots were probed with anti-GFP to detect the presence of HspB8 in the IP lysates or with **F)** anti-stv to detect the presence of stv in the IP lysates. **G)** Muscle lysates expressing control, UAS-mito-GFP, endogenously tagged stv or HspB8, or over-expressing MHC-GAL4 UAS-Lamin^GFP^ were immunoprecipitated using anti-GFP coated beads. Endogenous Lamin (~75kDa) was pulled down by stv-GFP. In addition, Lamin-GFP (~100kDa) was also detected as a control. Endogenous phoshop-p38K was pulled down by both stv and lamin. Endogenous stv (~80kDa) was pulled down by both HspB8 and Lamin. In addition, as a control we can detect pull down of the stv-GFP (~100kDa). Anti-GFP was used to confirm to the pull down of both GFP alone and HspB8-GFP.

### p38Kb physically interacts with the CASA complex in the adult flight muscle

In order to further determine if p38Kb physically interacts with the CASA complex in the adult muscle, we performed co-immunoprecipitation experiments. In *Drosophila* cell culture p38Kb^wt^ was shown to co-immunoprecipitate with the HspB8 homologue CG14207^18^, therefore, we tested if this interaction also occurs in adult indirect flight muscle. As the interaction between p38Kb and its targets may be transient, we utilized a FLAG-tagged p38Kb kinase dead construct (UAS-p38Kb^KD^ Mef2-GAL4) that is able to be activated and bind to a target but cannot phosphorylate it, leading to a delayed release of the target^36^. We expressed p38Kb^KD^ in the adult flight muscle of a fly that also endogenously expressed GFP tagged HspB8, in which GFP is spliced in as a new exon of the endogenous gene. This results in a GFP fusion protein that is expressed in the same pattern as endogenous HspB8 (^19^ and Figure 3C’). We find that p38Kb^KD^ was able to pull down HspB8 (Figure 3E) as compared to the HspB8-GFP control background. In addition, expression of p38Kb^KD^ in a wildtype background was able to pull down un-tagged endogenous stv (Figure 3F). We then performed the reverse immunoprecipitation experiments in which we immunoprecipitated control GFP or endogenously GFP tagged Hsc70-4, HspB8 or stv from muscle lysates and probed for endogenous phosphorylated p38K, which is the active form of p38K. We find that all three members of the CASA can co-immunoprecipitate with endogenous phosphorylated p38K in the muscle (Figure 3G, Figure S1D). We repeated these experiments in which we expressed the FLAG tagged p38Kb^KD^ and found that p38Kb co-immunoprecipitates with the CASA complex (Figure S1A-C) with stronger binding at younger ages than older ages. To determine if this reduction in binding was due to an age-dependent decrease in the ability of p38K to associate with the CASA complex, we tested if the expression levels of either the CASA complex or the FLAG tagged p38Kb^KD^ are changed with age. We found no significant difference in the expression levels of Hsc70-GFP and stv-GFP with age (Figure S2A and C) while levels of HspB8-GFP increased with age (Figure S2B). In addition, the levels of over-expressed p38Kb^KD^ declined with age (Figure S2D). We also have found that levels of p38K decline with age^37^. These data suggest that the levels of p38Kb protein may be limiting for CASA complex functions as the fly ages.

### p38Kb acts downstream of the CASA complex

We next tested the relationship between p38Kb and the CASA complex to determine if p38Kb acts either upstream or downstream of the CASA complex to promote protein homeostasis. As BAG-3/stv provides both the specificity to the CASA complex and is involved in the handoff of damaged proteins to p62/ref(2)p for autosomal degradation ^21–25^, we tested for genetic interactions between p38Kb and *stv*. *stv* null mutants have impaired locomotor functions, muscle degeneration and early lethality ^19,38^. Due to the severity of these null phenotypes, we utilized *stv* RNAi lines to generate an allelic series of *stv* loss of function in the muscle. We find that weak inhibition of stv (UAS-stv RNAi^34408^ MHC-GAL4) had no effect on protein aggregation (Table S6). However, moderate inhibition of stv (UAS-stv RNAi^34409^ MHC-GAL4) results in increased protein aggregate number and size (Figure 4A-B and Table S7) and results in a decrease in lifespan particularly in the first half of life as compared to outcrossed controls (Figure 4C and Table S8). Strong inhibition of *stv* (UAS-stv RNAi^34408^ Mef2-GAL4) leads to a severely reduced lifespan of ~4 days on average (Figure 4D and Table S9).

**Fig 4.**
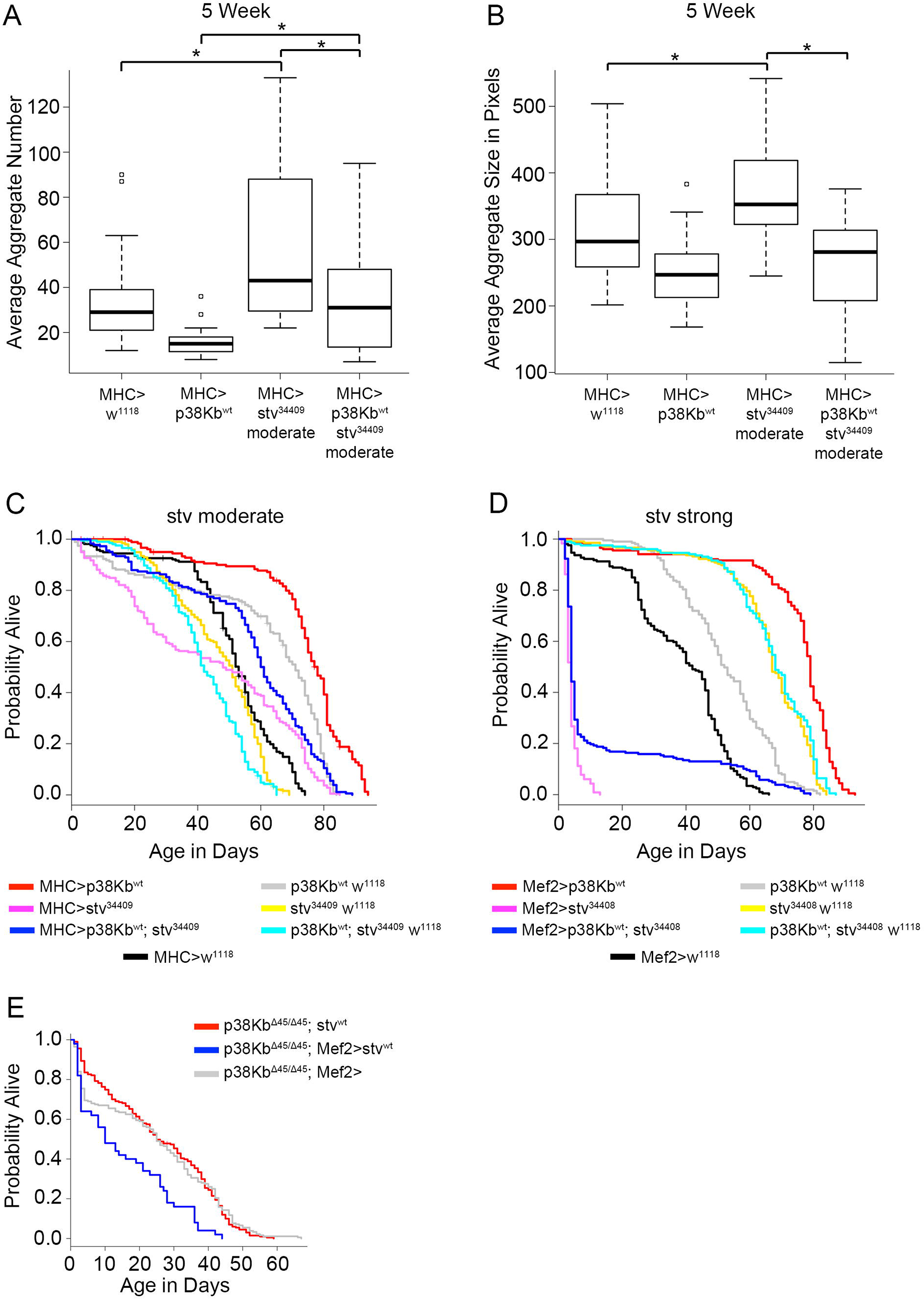
p38Kb genetically interacts with stv to regulate protein homeostasis and lifespan. **A)** Protein aggregate number and **B)** size in the moderate stv knockdown background using MHC-GAL4 at 5 weeks. Asterisks denote a p-value of ≤0.001. **C)** Moderate over-expression of p38Kb (red line) results in an increased lifespan as compared to the MHC-GAL4 controls and p38Kb transgene control (black line and gray lines, respectively). Moderate knockdown of stv using the MHC-GAL4 results in a decreased lifespan (pink line compared to yellow and black lines) and is rescued by p38Kb over-expression (compare pink line to blue line). **D)** Strong over-expression of p38Kb (red line) results in an increased lifespan as compared to the Mef2-GAL4 controls and p38Kb transgene control (black line and gray lines, respectively). Strong knockdown of stv using the Mef2-GAL4 results in a decreased lifespan (pink line compared to yellow and black lines) and is partially rescued by p38Kb over-expression (compare pink line to blue line). **E)** Over-expression of stv in the p38Kb mutant background results in a further reduction of lifespan as compared to p38Kb mutant controls (compare blue line to red and grey lines).

If p38Kb acts to regulate the downstream activity of the CASA complex, then over-expression of p38Kb should rescue these *stv* RNAi phenotypes. We find that over-expression of p38Kb rescues both protein aggregate number and size in the *stv* moderate inhibition background (Figure 4A-B and Table S7) and also rescues the lifespan defect (Figure 4C and Table S8). Furthermore, p38Kb over-expression is able to partially rescue the decreased lifespan exhibited by strong *stv* inhibition increasing average lifespan from 4 days to 13 days (Figure 4D and Table S9). To further test if p38Kb acts downstream of the CASA complex, we combined loss of p38Kb with CASA complex over-expression. If p38Kb acts downstream of the CASA complex, then over-expression of the CASA complex members in the p38Kb mutant background will not be able to rescue the p38Kb mutant phenotypes. We find that over-expression of Hsc70-4 in the muscle fails to rescue the p38Kb mutant shortened lifespan (Figure S3A, Table S10), even though this extends lifespan in a wild type background (Figure S3B, Table S11). Furthermore, we find that over-expression of stv in the p38Kb mutants not only fails to rescue the p38Kb mediated short lifespan defect but reduces the lifespan further (Figure 4E and Table S12). This is particularly striking as over-expression of stv in a wild type background does not significantly affect lifespan (Figure 5E and Table S13) as compared to the outcrossed transgene control. These data in combination with the decrease of p38K binding to the CASA complex with age when p38K levels are reduced suggest that p38Kb may be a limiting factor in regulating the downstream activity of the CASA complex in which it hands the poly-ubiquitinated misfolded proteins to ref(2)p for degradation through the autophagosome.

**Fig 5.**
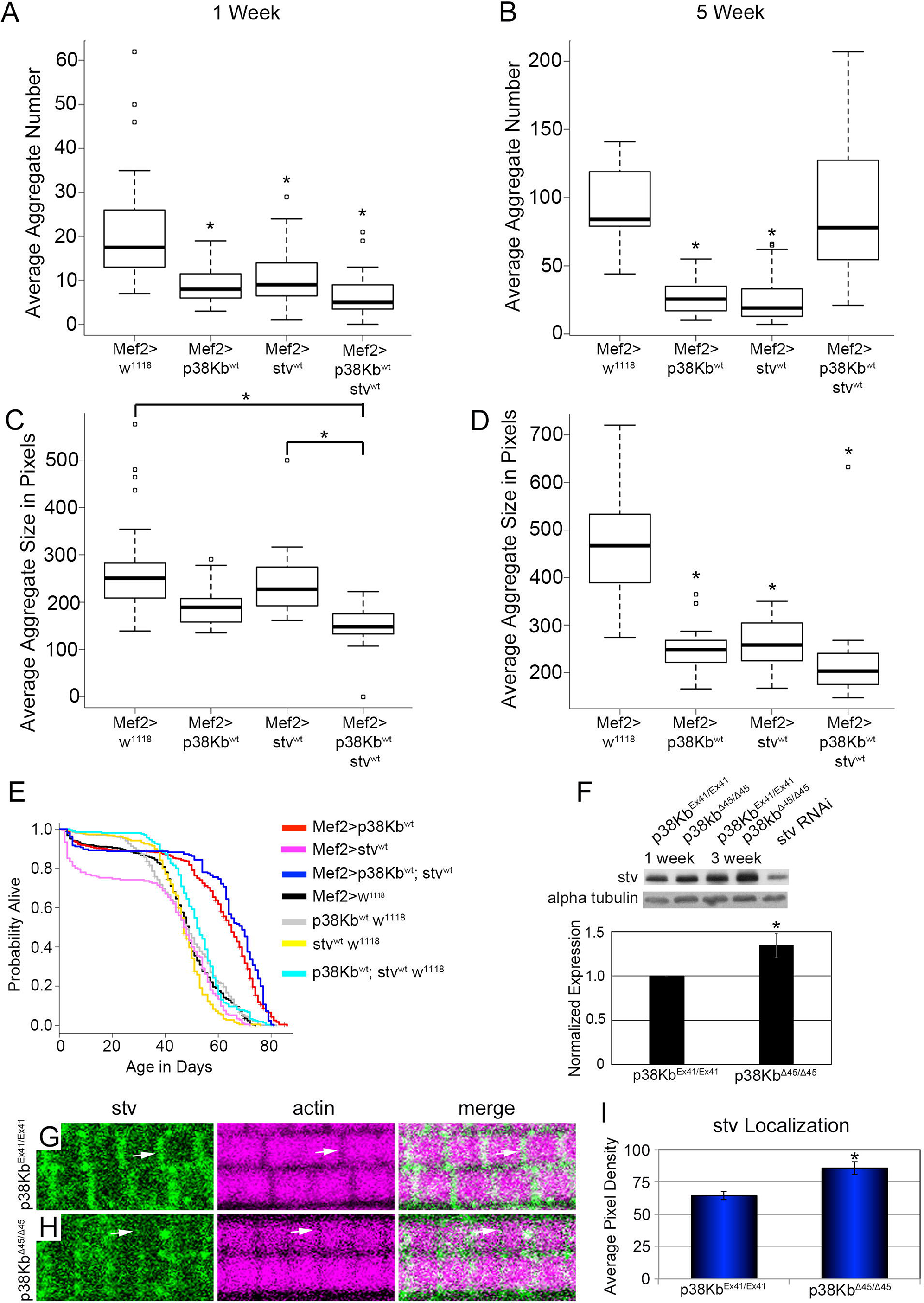
p38Kb is limiting for stv function and localization. Protein aggregate number at **A)** 1 week and **B)** 5 weeks and protein aggregate size at **C)** 1 week and **D)** 5 weeks measured in stv over-expression backgrounds. Over-expression of stv leads to reduced aggregate number at 1 and 5 weeks and aggregate size at 5 weeks. Co-over-expression of stv and p38Kb does not result in a further decrease in protein aggregate number but trends towards decreased aggregate size at both 1 and 5 weeks of age. Asterisks denote a p-value of ≤0.001 when compared to the GAL4 control. **E)** Over-expression of stv alone has no significant effect on lifespan (pink line as compared to yellow and black lines), however, co-over-expression of stv and p38Kb results in a further increase in lifespan (compare red line to blue line). **F)** Immunoblots of stv from 1 and 3 week old muscle lysates of control and p38Kb mutants. **G)** stv localizes to the adult muscle Z-disc and M-line (white arrows) in control animals. **H)** stv localization is disrupted in p38Kb mutants. **I)** Quantification of average pixel density.

### p38Kb regulates the activity of the CASA complex in protein homeostasis

If p38Kb is a limiting factor for stv function, then the combined over-expression of p38Kb and stv may result in a further beneficial effect. We find that over-expression of stv alone results in fewer aggregates at young and old ages (Figure 5A-B and Table S14) and smaller aggregates with age (Figure 5D and Table S14) as compared to outcrossed controls. However, co-over-expression of p38Kb and stv does not result in a further reduction in aggregate number as compared to over-expression of p38Kb or stv alone (Figure 5A and Table S14). By 5 weeks of age, co-over-expression of p38Kb and stv flies have comparable aggregate number to controls (Figure 5B and Table S14). Conversely, p38Kb and stv co-over-expression results in a reduction in aggregate size at a young age compared to over-expression of stv alone (Figure 5C and Table S14), suggesting that p38Kb is a limiting factor for stv function in regulating aggregate size. Unlike with aggregate number, the aggregates remain significantly smaller in size as the flies age in the combined over-expression background (Figure 5D and Table S14). We also find that co-over-expression of p38Kb and stv leads to an additional 5% increase in lifespan relative to p38Kb over-expression alone (Figure 5E and Table S13).

Interestingly, the co-over-expression animals show a very similar lifespan to the p38Kb over-expression alone animals until ~day 50, when the p38Kb alone animals begin to die at a faster rate (Figure 5E). These data suggest that co-over-expression of p38Kb and stv provides beneficial effects in early adulthood that continue throughout adulthood leading to increased lifespan despite the presence of protein aggregates. Another possibility is that aggregate size and/or content may play a more important role in determining lifespan as compared to overall aggregate number.

### p38Kb is required for proper localization of stv in the muscle

Since we find that p38Kb acts between stv and ref(2)p in regulating protein aggregation, we hypothesize that p38Kb plays a role in stabilizing the transfer of misfolded proteins primed for degradation from the CASA complex to ref(2)p. In support of this hypothesis, we find that over-expression of p38Kb leads to increased efficiency in targeting misfolded proteins to the autophagosome (Figure 1M-T), while inhibition of p38Kb results in increased protein aggregation (Figure 1E-L). In addition, loss of p38Kb may lead to a failure of stv to maintain its interaction with the CASA complex or to interact with ref(2)p. Therefore, we tested if stv is still able to localize to the Z-disc in the absence of p38Kb. We find that in p38Kb null mutants, stv protein levels are increased (Figure 5F), and stv is more diffusely located throughout the muscle but is still able to localize to the Z-disc and M-line in (Figure 5G-I), suggesting that its localization is partially impaired in the absence of p38Kb. Furthermore, we find that the localization of the CASA complex member HspB8 in the muscle is unaffected by loss of p38Kb (Figure S3C-D). These data indicate that p38Kb may play a role in stabilizing the interaction between the CASA complex and stv that allows stv to direct misfolded proteins to ref(2)p.

### Lamin protein binds to the CASA complex and accumulates in stv RNAi and p38Kb mutants

In order to determine if p38Kb might be playing a role in the targeting of misfolded proteins from stv to ref(2)p, we first needed to identify a protein target of both stv and p38Kb. Therefore, we focused on proteins associated with Limb-Girdle Muscular Dystrophy (LGMD), which is caused by mutations in either BAG-3 (stv, myofibrillar myopathy-6) or HspB8 (LGMD Type 1D) ^39–41^. In addition, p38K signaling has been implicated in LGMD ^42–44^. *Drosophila* have 19 orthologues of LGMD proteins, one of which is Lamin A/C (Lamin or Lamin Dm_0_ in flies). Lamin is of particular interest since mutations in Lamin A/C also result in the accelerated aging disorder Hutchinson-Gilford progeria ^45–47^, and Lamin has been shown to aggregate under oxidative stress conditions^48^. Mutations in Lamin A/C are also sufficient to induce Lamin aggregation and abnormal nuclear morphology in human cell culture, *C. elegans*, and *Drosophila* systems ^46,49–57^. In addition, Lamin proteins are highly post-translationally modified, resulting in changes in its solubility^56,58–63^. Furthermore, in *Drosophila* inhibition of either fly homologue of Lamin A/C have similar phenotypes to p38Kb and/or stv mutants, such as reduced locomotor function and increased activity of the Nrf-2/Keap-1 pathway ^51,64,65,57,66^. In addition, it has recently been shown that BAG-3, one of the mammalian homologues of stv, can target nuclear Lamin B, a paralogue of Lamin A/C, for degradation ^67^.

If Lamin is a target of the CASA complex, then decreased activity of the CASA complex should result in an accumulation of Lamin protein. Flies have two homologues of Lamin A/C, Lamin Dm_0_ and LamC. We find that the levels of Lamin Dm_0_, do not change with age (Figure 6A), however, inhibition of stv results in a significant increase in the total amount of Lamin Dm_0_ protein regardless of age (Figure 6A-B). Interestingly, loss of p38Kb leads to an age-dependent increase in Lamin Dm_0_ protein as compared to age matched controls (Figure 6C-D). These data in addition to our findings that p38Kb is limiting for stv localization (Figure 5H) and that the levels of p38Kb decline with age^37^ suggest that p38Kb activity later in life is important for maintaining protein homeostasis and lifespan.

**Fig 6.**
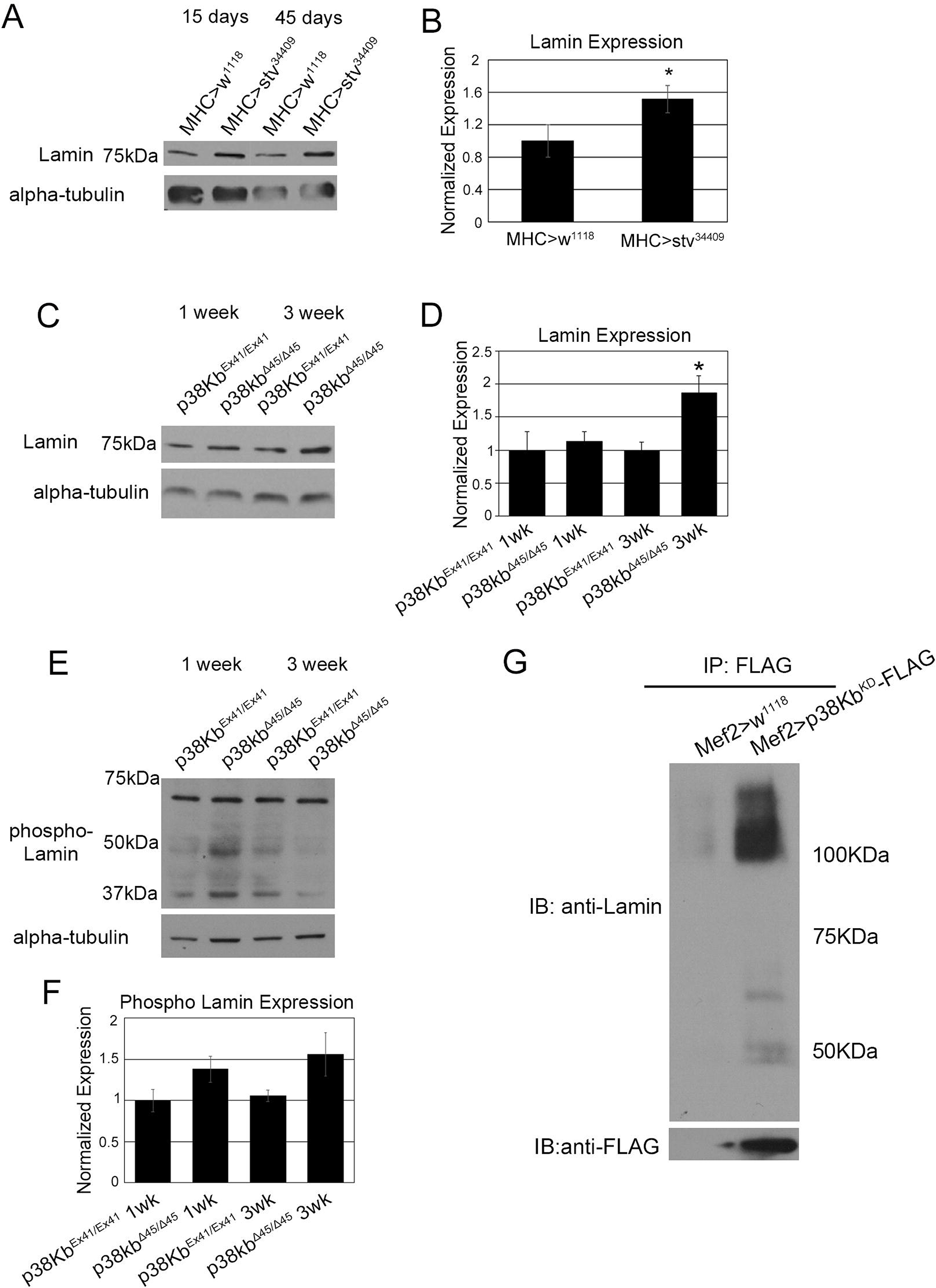
p38Kb and stv regulate Lamin aggregation. **A)** Immunoblot analysis of stv-RNAi and GAL4 controls flies muscle lysates probed with anti-Lamin and **B)** quantified using densitometry, asterisk denotes p value=0.028. **C)** Immunoblot analysis of p38Kb^Ex41/Ex41^ control and p38Kb ^Δ45/Δ45^ mutant muscle lysates probed with anti-Lamin and **D)** quantified using densitometry, asterisk denotes p value = 0.006643. **E)** Immunoblot analysis of p38Kb^Ex41/Ex41^ control and p38Kb ^Δ45/Δ45^ mutant muscle lysates probed with anti-phospho Lamin and **F)** quantified using densitometry, p=0.09. **G)** p38Kb^KD^ was expressed in muscles using Mef2-GAL4. p38Kb was immunoprecipitated from adult muscle lysates using anti-FLAG coated beads. Immunoblots were probed with anti-lamin and anti-FLAG.

To investigate if Lamin Dm_0_ is accumulating in the protein aggregates, we performed fractionation experiments in which we find that in controls Lamin Dm_0_ is predominantly found in fractions 5-7 and also in the aggregate containing pellet (Figure 7A), which is characterized by an increased amount of poly-ubiquitinate proteins (Figure 7C). Interestingly, we also find a low molecular weight from of Lamin Dm_0_ mainly restricted to fraction 5 and a higher molecular weight form that is predominantly in the pellet (Figure 7A). We next tested how loss of p38Kb affects the aggregation of Lamin Dm_0_ and find that p38Kb mutants have decreased Lamin Dm_0_ in fractions 5-7 with a concurrent increase of both the main and high molecular weight species of Lamin Dm_0_ in the pellet (Figure 7A). This suggests that Lamin Dm_0_ is a target of p38Kb and that loss of p38Kb results in increased aggregation of Lamin Dm_0_.

**Fig 7.**
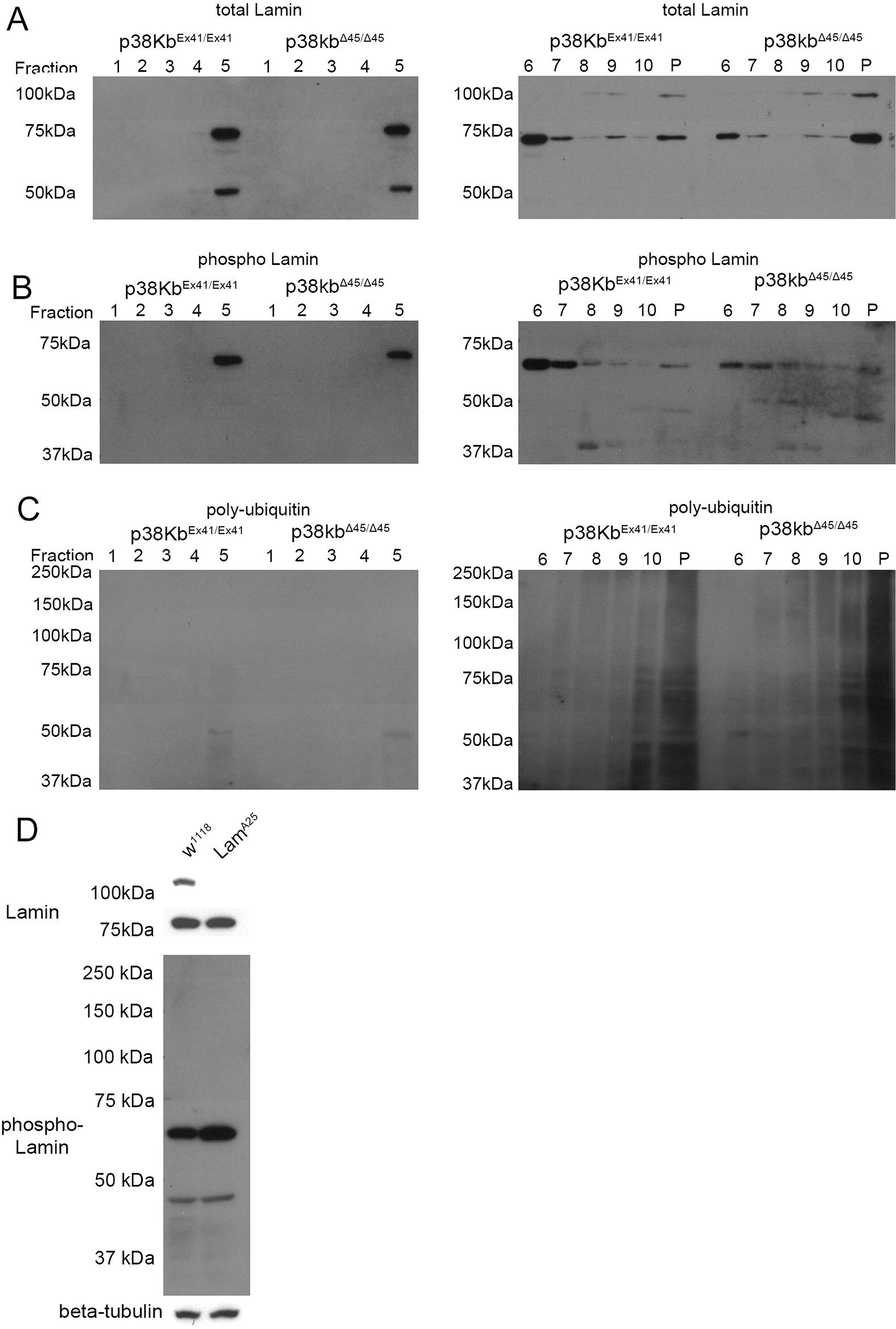
Lamin aggregation is increased in aged p38Kb mutants. **A-C)** Immunoblots of sucrose gradient fractions from 3 week old p38Kb^Ex41/Ex41^ control and p38Kb ^Δ45/Δ45^ mutant muscle. **A)** The main species of Lamin (~75kDa) is mainly found in fractions 5-7 with some in the pellet, and the farnesylated form of Lamin (~100kDa) is mostly found in the pellet in controls. In the p38Kb mutants, there is a further accumulation of Lamin and farnesylated Lamin in the pellet. **B)** Phosphorylated Lamin is also found predominantly in fractions 5-7 in controls and accumulates in the pellet in p38Kb mutants. **C)** Poly-ubiquitin is found mostly in the pellet fraction, marking the protein aggregates. **D)** Immunoblots of control and Lamin^A25^ mutants that lack the CAAX motif demonstrate the loss of the farnesylated form of Lamin (100kDa) without affecting the main form of Lamin (~75kDa). In addition, phosphorylated Lamin is also present in the Lamin mutants.

Of interest is this high molecular weight form of Lamin Dm_0_, which is present in the aggregates (Figure 7A). Lamin proteins are highly post-translationally modified^56,58–63^, and one important protein domain in Lamins is the CaaX box, which is farnesylated and required for Lamin localization to the inner nuclear membrane^63,68–70^. To test if this might be a farnesylated form of Lamin Dm_0_, we utilized the Lam^A25^ mutant which has a frameshift that results in the loss of the C-terminal CaaX box^71^. We find that the high molecular weight form of Lamin Dm_0_ is lost in the Lam^A25^ mutant whereas the predominant 75kDa band of Lamin Dm_0_ is still present (Figure 7D). These data suggest that the high molecular weight Lamin Dm_0_ we observe is the farnesylated form. We do not observe a consistent effect on levels due to either stv inhibition or loss of p38Kb, which suggest that p38Kb mediates farnesylated Lamin Dm_0_ aggregation than overall levels.

In addition to farnesylation, Lamins can also be phosphorylated by a variety of kinases that can change the solubility of Lamin porteins^56,72^. For example, in a *Drosophila* cell culture system, a phospho-mimic mutation at Ser45 (the equivalent of Ser22 in humans) results in Lamin aggregation in both the nucleus and cytoplasm ^56^. This serine residue is phosphorylated by the cd2 kinase ^73^, but is also a predicted p38K phosphorylation site. As the antibody epitope is conserved in both Lam Dm_0_ and LamC, we may not be able to detect specific changes in phosphorylated Lam Dm_0_ versus LamC, however, we can assess changes in the levels of Lamin phosphorylation in general. We find that phospho-Lamin levels are not significantly altered in the p38Kb mutants (Figure 6E-F), suggesting that p38Kb is not required for Ser45 phosphorylation. Furthermore, this phosphorylated form of Lamin runs at lower molecular weight than the main form of Lamin (Figure 7D top and middle panels), suggesting that Lamins in adult *Drosophila* muscle is further processed. As this phosphorylation site is near the epitope recognized by the total lamin antibody, it maybe that lamin phosphorylation obscures the ability of this antibody to detect this form of Lamin.

We next examined if phosphorylated Lamin is also present in the aggregates. We find that in controls, phosphorylated Lamin is present in factions 5-7 with low amounts in the pellet (Figure 7B). We also observe a low molecular weight phospho-Lamin form that appears in the pellet (Figure 7B). In the p38Kb mutants, phospho-Lamin expression is reduced in fractions 6 and 7 and accumulates in the pellet (Figure 7B). Furthermore, increasingly smaller low molecular weight species of phospho-Lamin are present in the p38Kb^Δ45^ mutants including the pellet (Figure 7B), suggesting that loss of p38Kb prevents the effective clearance of these Lamin cleavage forms. Interestingly, loss of the Lamin CaaX box does not affect the formation of these lower molecular weight forms, however, it does result in an increase in the phosphorylation of full-length Lamin (Figure 7D). Additionally, we do not detect phosphorylation positive high molecular weight forms of Lamin (Figure 7D), suggesting that farnesylated Lamin is not phosphorylated at Ser45. Overall these data suggest that Lamin is processed in a variety of different ways in the adult muscle and these different forms are prone to aggregation in the absence of p38Kb.

To determine if Lamin Dm_0_ is a direct target of p38Kb and the CASA complex, we tested if Lamin Dm_0_ can bind to the CASA complex. We immunoprecipitated endogenously GFP tagged HspB8, Hsc70-4 and stv from adult *Drosophila* muscle and probed for endogenous Lamin Dm_0_. We find that full-length Lamin Dm_0_ co-immunoprecipitates with all three members of the CASA complex (Figure 3G and Figure S1E), but do not detect binding with either the higher or lower molecular weight forms of Lamin Dm_0_ found in the fractions (Figure 7A and B). Furthermore, we expressed Lamin Dm_0_ tagged with GFP in the muscle and found that endogenous stv co-immunoprecipitates with Lamin Dm_0_ (Figure 3G and Figure S1F). These data suggest that the CASA complex is able to bind to full-length Lamin Dm_0_ for targeting to the autophagosome.

As we hypothesize that p38Kb is interacting with the CASA complex at the point in which the poly-ubiquitinated misfolded targets are handed over to ref(2)p for degradation, we tested if p38Kb also binds to Lamin Dm_0_. We find that p38Kb^KD^ expressed in the muscle co-immunoprecipitates with both high and low molecular weight forms of Lamin Dm_0_ (Figure 6G). Interestingly, p38Kb physically interacts with the aggregate prone low molecular weight species of Lamin Dm_0_ as opposed to the predominant full-length form of Lamin Dm_0_ (Figure 6G). More striking is that p38Kb^KD^ physically interacts with increasingly larger forms of Lamin Dm_0_ (Figure 6G), suggesting that p38Kb is interacting with the poly-ubiquitinated forms of Lamin Dm_0_. Furthermore, we have expressed GFP-tagged Lamin Dm_0_ in the muscle and were able to pull down phosphorylated p38K (Figure 3G). These data indicate that p38Kb activation may be important for the hand off of misfolded proteins that are tagged for lysosomal degradation from stv to ref(2)p.

We have developed a model in which misfolded proteins such as the main form of Lamin Dm_0_ are targeted by stv to the CASA complex. Those proteins that cannot be refolded are then tagged with poly-ubiquitin to signal their degradation through the autophagosome/lysosome pathway. These poly-ubiquitinated proteins are handed off by stv to ref(2)p in a process mediated by activated p38Kb. We hypothesize that this hand off is a rapid process as we were unable to detect the higher molecular weight poly-ubiquitinated forms binding to the CASA complex and were only able to capture these transient interactions using the p38Kb^KD^ construct. This model explains our finding that in p38Kb mutants these poly-ubiquitinated proteins accumulate and then form aggregates, and how over-expression of p38Kb would lead to increased efficiency of the poly-ubiquitinated proteins being targeted to the autophagosome. In agreement with our model, full-length Lamin Dm_0_ would bind to the CASA complex for refolding and release, or for poly-ubiquitination and transfer to ref(2)p for degradation through an interaction with p38Kb.

## Discussion

We find that the aging gene p38Kb regulates age-dependent protein homeostasis through an interaction with the CASA complex and is acting at a step between the poly-ubiquitination of an un-foldable target and its transfer to ref(2)p for autolysosomal mediated degradation. Interestingly, we find that p38Kb is important for the proper localization of stv to the Z-disc but does not affect the localization of other complex members. This suggests that p38Kb may play a role in maintaining the interaction between stv and the CASA complex which may be necessary for target transfer to ref(2)p. stv has a conserved MAPK docking site as well as eight potential p38K phosphorylation sites. Furthermore, the mammalian ref(2)p homolog p62 has been shown in vitro to bind to mammalian p38K through two domains ^74^, which are partially conserved in flies. Therefore, one possibility is that p38Kb mediated phosphorylation of stv facilitates the localization of stv to form a functional CASA complex at the Z-disc, where damaged proteins are rapidly turned over. Another possibility is that p38Kb binds to stv and ref(2)p, and that the phosphorylation of stv is required for target hand-over to ref(2)p so that in the absence of p38Kb, stv cannot transfer targets to ref(2)p. A consequence of this may be that stv and the protein target together are released from the CASA complex, leading to mislocalization of stv. As we don’t detect a decrease in phospho-Lamin in the p38Kb mutants or find that the p38Kb can pull-down phospho-Lamin, it is unlikely that p38Kb is directly phosphorylating the target proteins as a part of the stv-ref(2)p hand-over process.

How protein aggregation contributes to aging and disease has been an area of great interest. One outstanding question is if protein aggregation is a consequence or cause of aging. It has been hypothesized that protein aggregates accumulate with age as the amount of damaged or misfolded proteins increase. However, it is not clear whether or not these aggregating proteins are toxic leading to tissue dysfunction and a disease state. Previous studies have found that long-lived fly strains such as over-expression of Foxo or parkin result in reduced protein aggregate formation ^4,75^. Therefore, we hypothesized that decreased protein aggregation would lead to a lifespan extension, while increased protein aggregation would lead to a reduced lifespan. As expected, we find that the short-lived p38Kb mutants, which exhibit premature locomotor behavior defects^16^, have large and numerous protein aggregates. We also find that over-expression of p38Kb leads to decreased protein aggregation and increased lifespan.

However, while inhibition of stv results in increased aggregation and decreased lifespan, over-expression of stv leads to decreased aggregate number without a concurrent lifespan extension. These data suggest that protein aggregation and lifespan may be separable processes. The CASA complex is important for maintaining lifespan as inhibition of stv results in a shorter lifespan. However, as stv over-expression on its own doesn’t extend lifespan, this suggests that lifespan extension may not entirely depend on the CASA complex or that increasing the activity of the CASA complex alone is not sufficient to drive lifespan. It has been hypothesized that the CASA complex is selective and only targets a subset of proteins for refolding/degradation. This selectivity is provided by BAG-3 in mammals and presumably stv in flies, though the selectivity of the complex has not been fully characterized in flies. This selectivity may explain why the CASA complex alone does not extend lifespan, as it may not target proteins that are known to cause increased lifespan but rather targets proteins that are involved in maintaining organismal health/lifespan.

Another potential interpretation of these results is that aggregate number is not as critical as aggregate size and/or content, particularly early in life, for lifespan extension. We find that stv over-expression only results in decreased aggregate size in older animals, however, co-over-expression of p38Kb and stv leads to reduced aggregate size at both young and old ages and leads to a further increase in lifespan. Thus, the turnover of specific toxic protein or subset of proteins early in life may lead to a reduction in the exposure to these toxic species and a lifespan extension. Furthermore, it has been hypothesized that protein aggregates may be protective in some instances. For example in Alzheimer’s disease it has been hypothesized that the soluble form of Amyloid-β is toxic and that the formation of aggregates protects against this toxicity ^76–80^. If aging or lifespan is dictated by the presence of soluble toxic proteins, then reducing these toxic proteins would increase lifespan. As over-expression of stv is unable to extend lifespan, this would then suggest that these particular soluble toxic proteins are not targets of the CASA complex.

Our data also suggest that the p38Kb can mediate lifespan through additional, CASA complex-independent mechanisms. Indeed, we have previously shown that p38Kb regulates lifespan at least in part by regulating oxidative stress^16^. Additionally, we find that over-expression of Hsc70-4, which is involved multiple protein quality control pathways, is able to extend life span, supporting the idea that CASA alone may not be sufficient to clear out all the toxic protein species that play a role in aging.

It has recently been found that over-expression of the related Hsp70 Ab gene in *Drosophila* adult muscle does not extend lifespan^81^. One possibility as to why we observe a lifespan extension is that we over-expressed Hsc70-4 using a different GAL4 driver (Mef2-GAL4 as compared to DJ694-GAL4) that drives expression all through development and in the adult though expression in the adult declines with age (Figure S2D). It may be that this early over-expression of Hsc70-4 provides a beneficial effect that protects the fly from general life stress, resulting in increased longevity. Another possibility is that Hsc70-4 and Hsp70 Ab proteins, which are 89.6% similar and 73% identical, have different cellular specificities that give rise to different functions resulting in different lifespan responses.

We find that Lamin, which is mutated to an aggregation prone protein form in Hutchinson-Gilford progeria ^46,55,57,72,82^, is a target of p38Kb and the CASA complex. Interestingly, p38Kb and Lamin mutants share similar phenotypes such as age-dependent locomotor impairments ^16,64,65^ and upregulation of the Nrf-2/Keap-1 pathway ^16,51^. As loss of p38Kb leads to an accumulation of Lamin in the aggregates, this may be toxic to the cell, leading to impaired locomotor function, increased stress and decreased lifespan. Thus, we have found that one aging gene (p38Kb) regulates a second, unrelated aging gene (Lamin) via the CASA complex. These data suggest a new link between aging pathways and how they may converge through the regulation of protein homeostasis.

## Acknowledgements

We would also like to acknowledge the Bloomington Drosophila Stock Center (NIH P40OD018537), the Vienna Drosophila Resource Center, and Kyoto Stock Center for providing fly stocks used in this study. A. Vrailas-Mortimer was funded by start-up funds from the University of Denver, a Knoebel Center for the Study of Aging pilot grant, start-up funds from Illinois State University and was supported by the National Institute of Arthritis and Musculoskeletal and Skin Diseases of the National Institutes of Health under Award Number R15AR070505. The content is solely the responsibility of the authors and does not necessarily represent the official views of the National Institutes of Health. S. Sanyal was funded by startup funds from Emory University. S. Ryan was supported by a Knoebel Center for the Study of Aging pilot grant to A. Vrailas-Mortimer and S. Barbee, N. Mortimer was funded by start-up funds from Illinois State University. The funders had no role in study design, data collection and analysis, decision to publish, or preparation of the manuscript.

---

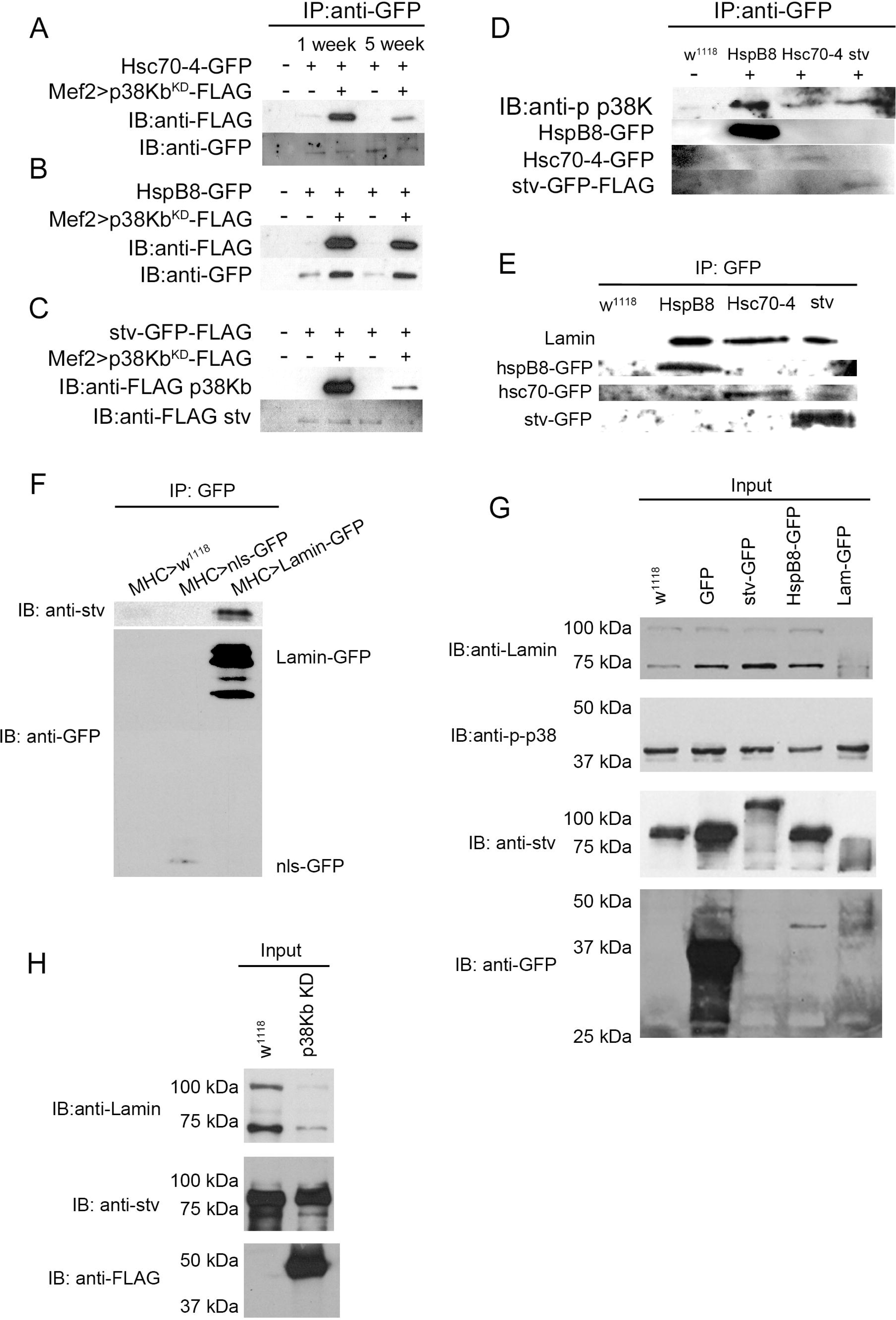

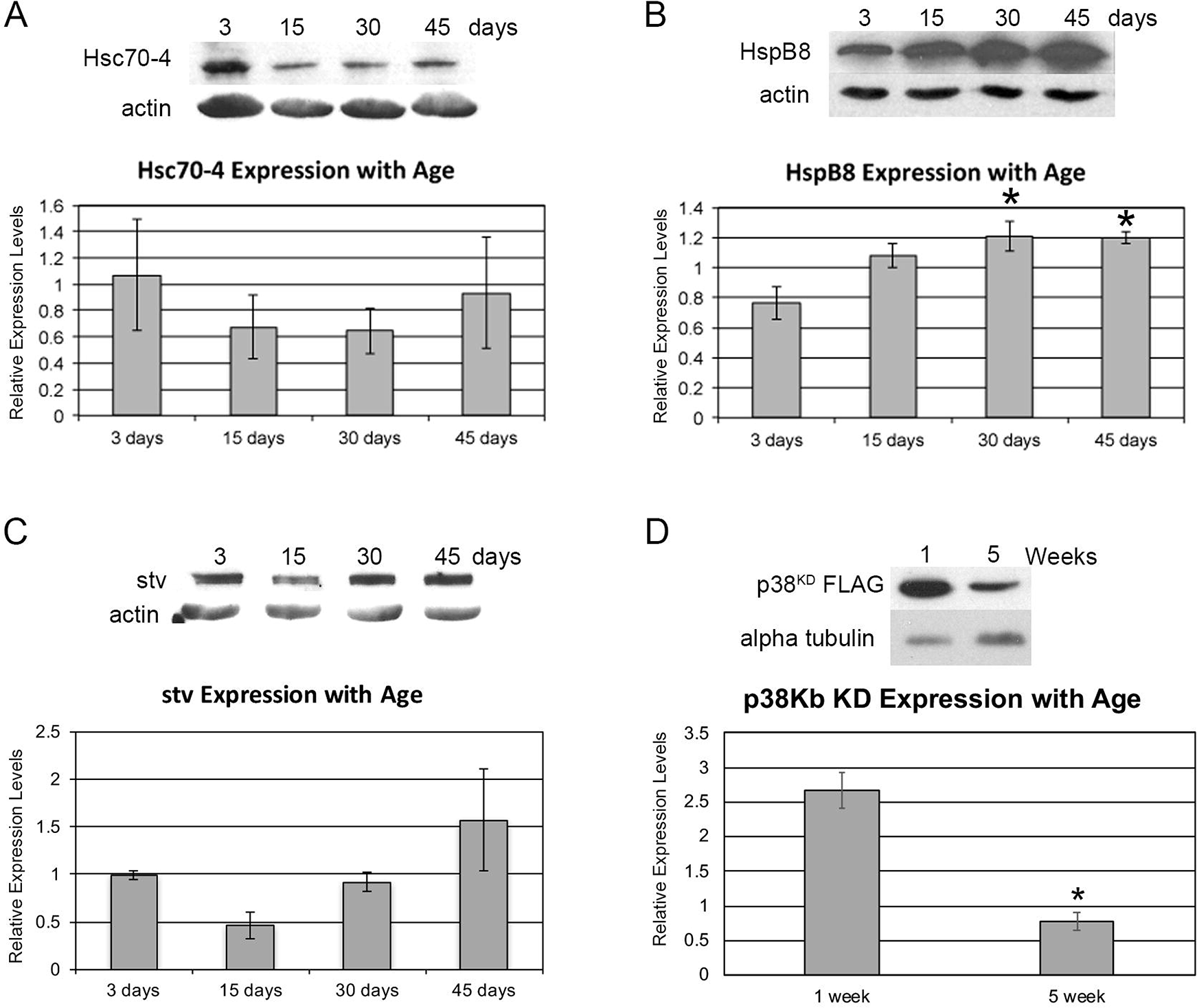

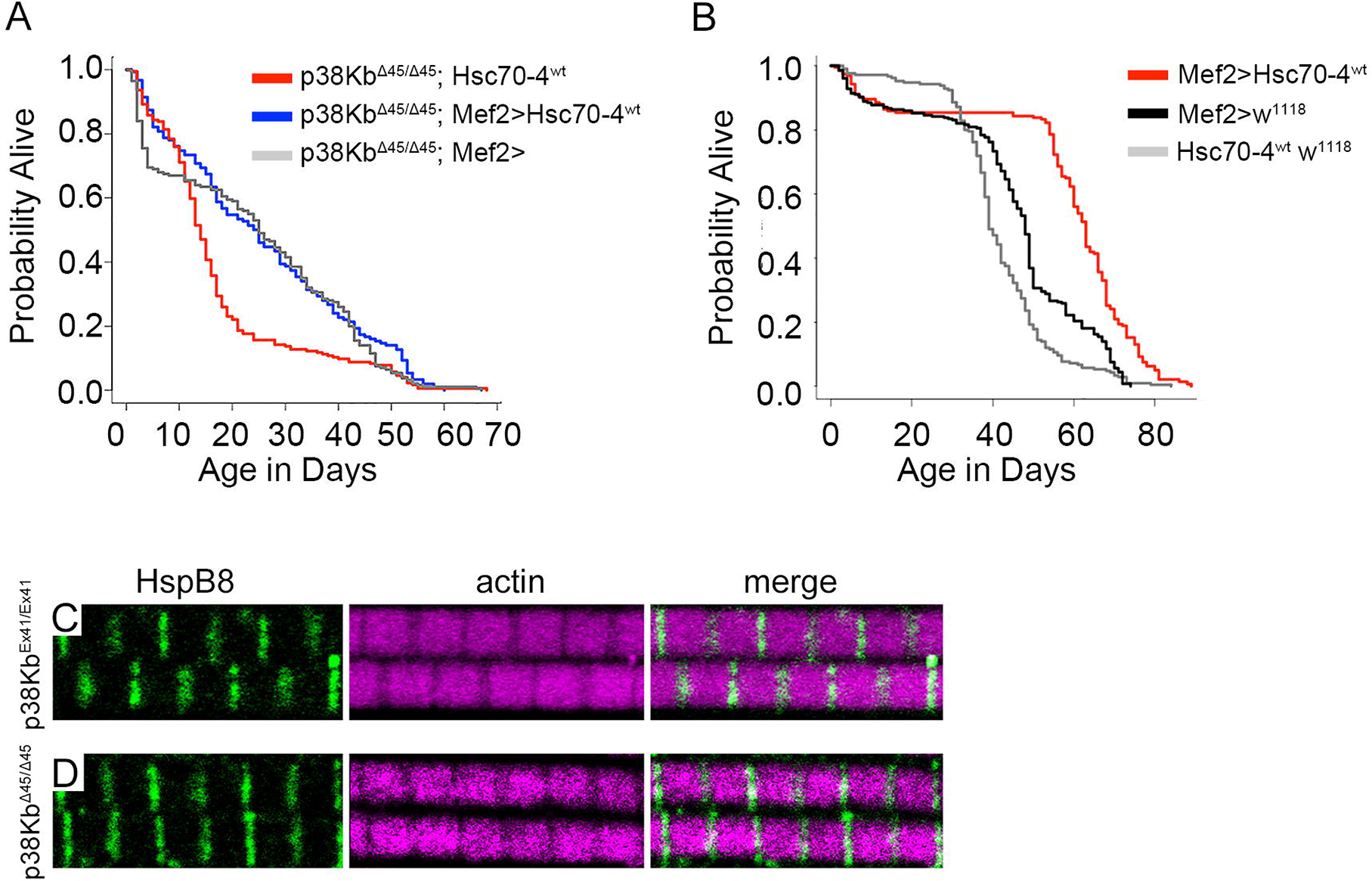

## Notes

### Competing Interest Statement

The authors have declared no competing interest.

### Summary of Updates

Figures have been revised with new data included.

