## Supplemental tables for "*Drosophila* p38 MAPK Interacts with BAG-3/starvin to Regulate Age-dependent Protein Homeostasis"

| Genotype | Aggregate Number 1 Week | Aggregate Number 3 Week | Aggregate Size 1 Week | Aggregate Size 3 Week |
| --- | --- | --- | --- | --- |
| p38Kb $^{Ex41/Ex41}$ | 8.8 ± 0.99 | 45.6 ± 5.09 | 209.8 ± 14.79 | 190.7 ± 8.15 |
| p38Kb $^{\Delta 45/\Delta 45}$ | 38.5 ± 4.28* | 139.19 ± 10.11* | 214.2 ± 14.38 | 385.2 ± 18.14* |

Table S1. Loss of p38Kb affects aggregate number and size. Mean aggregate number and size at 1 and 3 weeks. Asterisks indicate a p value of < 0.001

| Genotype | Aggregate Number 1 Week | Aggregate Number 5 Week | Aggregate Size 1 Week | Aggregate Size 5 Week |
|---|---|---|---|---|
| Mef2>$w^{1118}$ | 6.1 ± 1.18 | 57.6 ± 6.27 | 264.9 ± 21.09 | 540.09 ± 39.76 |
| Mef2> $p38Kb^{KD}$ | 16.7 ± 3.94* | 91.3 ± 11.67* | 277.3 ± 23.30 | 631.7 ± 48.92 |

Table S2. Expression of a dominant negative p38Kb Kinase Dead affects aggregate number. Mean aggregate number and size at 1 and 5 weeks. Asterisks indicate a p value of <0.02

| Genotype | Aggregate Number 1 Week | Aggregate Number 5 Week | Aggregate Size 1 Week | Aggregate Size 5 Week |
|---|---|---|---|---|
| Mef2>$w^{1118}$ | 24.5 ± 2.19 | 92.0 ± 3.72 | 310.3 ± 19.91 | 398.8 ± 13.69 |
| Mef2>$p38Kb^{wt}$ | 6.7 ± 0.47* | 37.7 ± 2.35* | 199.2 ± 6.81* | 287.1 ± 10.03* |

Table S3. Strong over-expression of p38Kb affects aggregate number and size. Mean aggregate number and size at 1 and 5 weeks. Asterisks indicate a p value of < 0.001

| Genotype | Aggregate Number 1 Week | Aggregate Number 5 Week | Aggregate Size 1 Week | Aggregate Size 5 Week |
|---|---|---|---|---|
| MHC>$w^{1118}$ | 18.0 ± 2.02 | 63.9 ± 5.83 | 332.3 ± 19.02 | 375.2 ± 15.74 |
| MHC>$p38Kb^{wt}$ | 5.8 ± 0.59* | 17.5 ± 1.52* | 219.8 ± 15.6* | 244.4 ± 14.40* |

Table S4. Moderate over-expression of p38Kb affects aggregate number and size. Mean aggregate number and size at 1 and 5 weeks. Asterisks indicate a p value of < 0.001

| Genotype | Aggregate Number 1 Week | Aggregate Number 5 Week | Aggregate Size 1 Week | Aggregate Size 5 Week |
|---|---|---|---|---|
| Mef2>w1118 | 12.6 ± 1.01 b | 100.0 ± 6.55 c | 248.0 ± 24.77 b | 334.1 ± 13.43 ab |
| Mef2>p38Kb$^{wt}$ | 4.0 ± 0.56 a | 34.9 ± 2.59 a | 174.8 ± 9.01 a | 381.1 ± 16.68 b |
| Mef2>ref(2)p$^{-/+}$ | 7.6 ± 1.15 ab | 74.2 ± 5.69 b | 181.6 ± 11.79 a | 324.1 ± 13.66 a |
| Mef2>p38Kb$^{wt}$ ref(2)p-/+ | 19.6 ± 2.46 c | 108.1 ± 7.83 c | 185.6 ± 5.93 a | 339.8 ± 13.86 ab |

Table S5. Reduction of ref(2)p affects p38Kb aggregate number. Mean aggregate number and size at 1 and 5 weeks. Means not sharing the same letter (a, b, c) are significantly different (Tukey's HSD, $p<0.05$).

| Genotype | Aggregate Number 1 Week | Aggregate Number 5 Week | Aggregate Size 1 Week | Aggregate Size 5 Week |
|---|---|---|---|---|
| MHC>$w^{1118}$ | 12.5 ± 1.21 b | 93.2 ± 7.43 b | 309.9 ± 19.15 a | 437.3 ± 22.23 c |
| MHC>$p38Kb^{wt}$ | 5.1 ± 0.48 a | 19.4 ± 2.81 a | 228.1 ± 24.89 a | 236.6 ± 27.39 a |
| MHC>$stv^{34408}$ RNAi | 11.4 ± 1.35 b | 72.1 ± 9.05 b | 293.3 ± 30.79 a | 362.0 ± 9.66 b |
| MHC>$p38Kb^{wt}$ $stv^{34408}$ RNAi | 14.1 ± 1.84 b | 29.8 ± 4.23 a | 233.9 ± 11.93 a | 261.7 ± 12.84 a |

Table S6. Mild inhibition of stv has no effect on p38Kb aggregate number or size. Mean aggregate number and size at 1 and 5 weeks. Means not sharing the same letter (a, b, c) are significantly different (Tukey's HSD, $p<0.05$).

| Genotype | Aggregate Number 1 Week | Aggregate Number 5 Week | Aggregate Size 1 Week | Aggregate Size 5 Week |
|---|---|---|---|---|
| MHC>$w^{1118}$ | 23.6 ± 3.58 bc | 33.6 ± 3.87 b | 354.6 ± 33.15 b | 310.8 ± 14.25 b |
| MHC>$p38Kb^{wt}$ | 6.4 ± 1.08 a | 15.6 ± 1.20 a | 210.5 ± 17.22 a | 252.2 ± 10.32 a |
| MHC>$stv^{34409}$ RNAi | 28.2 ± 6.40 c | 60.4 ± 6.99 c | 234.1 ± 15.28 a | 363.1 ± 13.67 c |
| MHC>$p38Kb^{wt}$ $stv^{34409}$ RNAi | 9.8 ± 1.75 ab | 35.5 ± 4.90 b | 281.7 ± 40.04 ab | 263.9 ± 13.62 ab |

Table S7. Moderate inhibition of stv has affects p38Kb aggregate number but not size. Mean aggregate number and size at 1 and 5 weeks. Means not sharing the same letter (a, b, c) are significantly different (Tukey’s HSD, $p<0.05$).

| Genotype | Average Age | Median Age | n | p value vs MHC>$stv^{34409}$ RNAi | p value vs MHC>$p38Kb^{wt}$ | p value vs MHC>$w^{1118}$ | p value vs Transgene Control |
|---|---|---|---|---|---|---|---|
| MHC>$w^{1118}$ | 51 days | 52 days | 212 | **4.00E-03** | 0 | - | - |
| MHC>$p38Kb^{wt}$ | 73.5 days | 77 days | 175 | **0** | - | 0 | 6.48E-13 |
| MHC>$stv^{34409}$ RNAi | 44.3 days | 48 days | 217 | - | **0** | **4.00E-03** | **3.32E-06** |
| MHC>$p38Kb^{wt}$ $stv^{34409}$ RNAi | 56.9 days | 60.5 days | 182 | **1.49E-04** | **0** | 3.54E-12 | 0 |
| $p38Kb^{wt}$ $w^{1118}$ | 59.4 days | 70 days | 178 | 1.93E-09 | 6.48E-13 | 0 | - |
| $stv^{34409}$ $w^{1118}$ | 46.2 days | 50.5 days | 194 | 3.31E-06 | 0 | 7.78E-06 | - |
| $p38Kb^{wt}$ $stv^{34409}$ RNAi $w^{1118}$ | 41.3 days | 41 days | 197 | 7.68E-10 | 0 | 0 | - |

Table S8. Moderate inhibition of stv prevents p38Kb lifespan extension. Chisq = 551, p = 0.

| **Genotype** | **Average Age** | **Median Age** | **n** | **p value vs Mef2>stv$^{34408}$** | **p value vs Mef2>p38Kb$^{wt}$** | **p value vs Mef2>w$^{1118}$** | **p value vs Transgene Control** |
|---|---|---|---|---|---|---|---|
| Mef2>w1118 | 38 days | 42 days | 206 | 0 | 0 | - | - |
| Mef2>p38Kb$^{wt}$ | 74.2 days | 79 days | 203 | 0 | - | 0 | 0 |
| Mef2>stv$^{34408}$ RNAi | 4.2 days | 4 days | 116 | - | **0** | **0** | **0** |
| Mef2>p38Kb$^{wt}$ stv$^{34408}$ RNAi | 13.4 days | 4 days | 208 | 1.86E-07 | **0** | 1.33E-10 | 0 |
| p38Kb$^{wt}$ w$^{1118}$ | 52.2 days | 52 days | 203 | 0 | 0 | 0 | - |
| stv$^{34408}$ RNAi w$^{1118}$ | 65.8 days | 67 days | 206 | 0 | 0 | 0 | - |
| p38Kb$^{wt}$ stv$^{34408}$ RNAi w$^{1118}$ | 66.3 days | 68 days | 203 | 0 | 6.48E-16 | 0 | - |

Table S9. Strong inhibition of stv prevents p38Kb lifespan extension. Chisq = 1693, p = 0.

| Genotype | Average Age | Median Age | n | p value vs GAL4 Control | p value vs transgene control |
|---|---|---|---|---|---|
| Mef2>p38Kb$^{\Delta45/\Delta45}$ | 24.14 days | 25 days | 200 | - | **0.00025** |
| p38Kb$^{\Delta45/\Delta45}$ Hsc70-4$^{wt}$ | 19.80 days | 15 days | 172 | **0.00025** | - |
| Mef2>p38Kb$^{\Delta45/\Delta45}$ ; Hsc70-4$^{wt}$ | 25.30 days | 24 days | 152 | 0.32214 | **9.50E-06** |

Table S10. Over-expression of Hsc70-4 fails to rescue p38Kb mutant short lifespan. Chisq= 25.1, p= 4e-06.

| Genotype | Average Age | Median Age | n | p value vs Mef2>$w^{1118}$ | p value vs Transgene Control |
|---|---|---|---|---|---|
| Mef2>w1118 | 44.5 days | 49 days | 278 | - | **2.10E-06** |
| Hsc70-$4^{wt}$ $w^{1118}$ | 41.4 days | 39 days | 210 | **2.10E-06** | - |
| Mef2>Hsc70-$4^{wt}$ | 56.32 days | 63 days | 192 | **< 2e-16** | **< 2e-16** |

Table S11. Hsc70-4 over-expression further extends lifespan. Chisq = 149, p < 2e-16.

| Genotype | Average Age | Median Age | n | p value vs GAL4 Control | p value vs transgene control |
|---|---|---|---|---|---|
| Mef2>p38Kb$^{\Delta45/\Delta45}$ | 24.1 days | 25 days | 200 | - | - |
| p38Kb$^{\Delta45/\Delta45}$ stv$^{wt}$ | 25.6 days | 25 days | 207 | 0.936 | - |
| Mef2>p38Kb$^{\Delta45/\Delta45}$ ; stv$^{wt}$ | 15.5 days | 10 days | 50 | **1.03E-04** | **1.87E-06** |

Table S12. Over-expression of stv fails to rescue p38Kb mutant short lifespan. Chisq = 22.7, p = 1.2E-05.

| Genotype | Average Age | Median Age | n | p value vs Mef2>stv$^{wt}$ | p value vs Mef2>p38Kb$^{wt}$ | p value vs Mef2>w$^{1118}$ | p value vs Transgene Control |
|---|---|---|---|---|---|---|---|
| Mef2>w1118 | 46.3 days | 49 days | 515 | 0.03458824 | 0 | - | - |
| Mef2>p38Kb$^{wt}$ | 58.2 days | 64 days | 730 | 0 | - | 0 | 0 |
| Mef2>stv$^{wt}$ | 39.7 days | 47 days | 182 | - | **0** | **0.034588** | 0.195 |
| Mef2>p38Kb$^{wt}$ stv$^{wt}$ | 61.3 days | 69 days | 205 | 0 | **0.03651667** | 0 | 0 |
| p38Kb$^{wt}$ w$^{1118}$ | 47.4 days | 48 days | 598 | 3.51E-04 | 0 | 0.1806 | - |
| stv$^{wt}$ w$^{1118}$ | 46.1 days | 47 days | 189 | 0.195 | 0 | 3.47E-03 | - |
| p38Kb$^{wt}$ stv$^{wt}$ w$^{1118}$ | 51.9 days | 52 days | 206 | 1.17E-04 | 0 | 2.00E-03 | - |

Table S13. p38Kb and stv co-over-expression further extends lifespan. Chisq = 706, p = 0.

| Genotype | Aggregate Number 1 Week | Aggregate Number 5 Week | Aggregate Size 1 Week | Aggregate Size 5 Week |
|---|---|---|---|---|
| Mef2>w1118 | 21.8 ± 2.64 b | 92.7 ± 5.10 b | 276.1 ± 21.07 c | 472.0 ± 20.95 b |
| Mef2>p38Kb$^{wt}$ | 8.7 ± 0.77 a | 27.5 ± 2.41 a | 192.2 ± 7.87 ab | 249.0 ± 8.11 a |
| Mef2>stv$^{wt}$ | 10.3 ± 1.22 a | 25.7 ± 3.39 a | 242.0 ± 13.51 bc | 258.6 ± 9.45 a |
| Mef2>p38Kb$^{wt}$ stv$^{wt}$ | 6.8 ± 1.01 a | 92.5 ± 9.48 b | 153.3 ± 5.68 a | 221.3 ± 17.29 a |

Table S14. Over-expression of stv affects p38Kb aggregate size at a young age. Mean aggregate number and size at 1 and 5 weeks. Means not sharing the same letter (a, b, c) are significantly different (Tukey's HSD, $p<0.05$).
